# Distinct regulation across the transcription cycle drives mitotic transcriptional silencing

**DOI:** 10.64898/2026.10.12.758451

**Authors:** Paola Gaetani, Alexandra Tavares, Tiago Paixão, Raquel A. Oliveira

## Abstract

In most organisms, as cells enter mitosis, transcription is globally silenced. While individual studies have shown that each stage of the transcription cycle can be modulated during mitosis, how initiation, elongation, and termination are temporally and kinetically regulated to ensure transcriptional silencing remains unclear. Here, we combine real-time quantitative imaging of ongoing transcription in early *Drosophila* embryos with mathematical modelling to systematically dissect how transcriptional kinetics change as cells progress into mitosis. We found that transcription initiation undergoes a gradual repression starting in prophase, while elongation rates remain unaltered. After nuclear envelope breakdown, an abrupt mechanism causes premature termination of elongating RNA, driven at least in part by the SNF-helicase-like protein Transcription Termination factor 2 / Lodestar (TTF2/Lds). Our study provides a quantitative framework for understanding how transcription is modulated across the cell cycle, demonstrating that mitotic repression arises from multiple processes with different dynamics that collectively ensure the precise shutdown of gene expression.

## Introduction

Multicellular organisms downregulate transcription during mitosis (Contreras & Perea-Resa, 2024; Edgar, 1986; Fink & Turnock, 1977; Gottesfeld & Forbes, 1997; Hartl et al., 1993; Prescott & Bender, 1962; Taylor, 1960). The process of transcription repression during mitosis is often intuitively attributed to the compaction of chromosomes upon mitotic entry. However, inhibiting or depleting the proteins responsible for mitotic chromosome organization (e.g. condensin I or topoisomerase 2) does not affect the extent or dynamics of transcription inhibition (Carmo et al., 2023; Hartl et al., 1993). On the contrary, it has been shown that chromosome accessibility does not change during mitosis (Ginno et al., 2018; Hsiung et al., 2015; Yu et al., 2023) and that several transcription factors remain bound to mitotic chromatin, a phenomenon known as mitotic bookmarking (Bellec et al., 2022; Burke et al., 2005; Caravaca et al., 2013; Christova & Oelgeschläger, 2002; Michelotti et al., 1997; Teves et al., 2016, 2018; Yang et al., 2013; Young et al., 2007). Furthermore, it has been proposed that some loci escape the general downregulation and are expressed during mitosis, giving this stage of the cell cycle its own expression profile (Palozola et al., 2017; Timmers & Verrijzer, 2017). Active mechanisms must therefore be in place to ensure prompt transcriptional repression.

Several studies highlighted how mitotic entry can have an impact on the different steps of the transcription cycle – initiation, elongation and termination. Several components of the basal transcriptional apparatus and transcription factors are hyperphosphorylated during mitosis (Akoulitchev & Reinberg, 1998; Cisek & Corden, 1989; Gebara et al., 1997; Hartl et al., 1993; Long et al., 1998; Lüscher & Eisenman, 1992; Rizkallah et al., 2011; Saad et al., 2025; Segil, 1996; Segil et al., 1991). The phosphorylation has been proposed to be mediated by the mitotic kinase Cdk1(Cisek & Corden, 1989; Gebara et al., 1997; Hartl et al., 1993; Leresche et al., 1996a; Long et al., 1998; Lüscher & Eisenman, 1992; Saad et al., 2025) and to negatively affect the DNA-binding activity of the target proteins (Lüscher & Eisenman, 1992; Martínez-Balbás et al., 1995; Rizkallah et al., 2011; Saad et al., 2025; Segil et al., 1991). While hyperphosphorylation has been suggested to prevent new transcription initiation events, the fate of already engaged polymerases is less clear. In HeLa cells, it has been shown that promoter escape is enhanced at mitotic onset to release RNA polymerases into elongation and clear them from gene bodies (Liang et al., 2015). In *Drosophila* embryos, entry into mitosis causes a progressive loss of nascent transcripts, which have been proposed to be prematurely terminated (Rothe et al., 1992; Shermoen & O’Farrell, 1991). These studies suggest that different steps of the transcription cycle can be regulated upon mitotic entry to ensure transcriptional silencing. Whether these multiple layers of regulation are simultaneously active within the same cell type, with which timings they act, and whether they crosstalk has not been determined yet.

The mechanisms that repress mitotic transcription must be particularly precise during early development. In many multicellular organisms, the first cell divisions after fertilization are extremely fast, with only a short window of time in which genes can be transcribed before being repressed at mitotic entry. Here, we used *Drosophila* syncytial embryos to understand how the stages of the transcription cycle are regulated as nuclei enter mitosis *in vivo*. At the syncytial stage of embryogenesis, the nuclei are arranged in a monolayer and divide synchronously every 13-15 min (Farrell & O’Farrell, 2014; Foe & Alberts, 1983). In addition, starting from the nuclear cycle 10 (nc10), the zygotic genome is partially active (Edgar, 1986; Farrell & O’Farrell, 2014; Pritchard & Schubiger, 1996; Zalokar, 1976). During this minor zygotic wave of gene expression, some genes are transcribed through a simplified transcription cycle, in which promoter-proximal pausing is abolished (Chen et al., 2013). Thus, the syncytial blastoderm represents an ideal system to study how transcription is regulated in nuclei approaching mitosis, as the zygotic genome is partially active and the cell divisions are fast, synchronous and accessible for imaging.

## Results

### Quantitative analysis of transcriptional output relative to cell cycle progression

To understand how the transcription cycle is affected as cells approach mitosis, we adopted a live imaging approach to achieve high temporal resolution during the fast embryonic divisions. Live imaging of transcription has been made possible using arrays of MS2 loops inserted in loci of interest. When the MS2 loops are transcribed, they are bound by MCP proteins tagged with a fluorophore, marking the nascent transcripts at the transcription site (Bertrand et al., 1998; Forrest & Gavis, 2003) (Figure 1A, B). For our analysis, we used an *MS2-lacZ-PP7* reporter under the control of the *hunchback* (*hb*) P2 promoter and enhancer (Fukaya et al., 2017) (Figure 1A). The *hunchback* promoter is active and shows no promoter-proximal pausing during the minor zygotic wave (Chen et al., 2013). We imaged embryos in nuclear cycle 12 (Figure 1B), and plotted the individual MCP-GFP signals, aligning them at the time of NEBD (Nuclear Envelope Break Down) (Figure 1C and S1). In agreement with previous reports (Carmo et al., 2023; Esposito et al., 2016; Garcia et al., 2013), we observed that the transcription signal decayed gradually upon mitotic entry (chromosome compaction above 10%, see Figure S2 for details on mitotic progression estimates) (Figure 1B-C and S1). However, imaging of total MCP signals cannot tell us how the steps of the transcription cycle – initiation, elongation, and termination - are affected during cell cycle progression.

**Figure 1.**
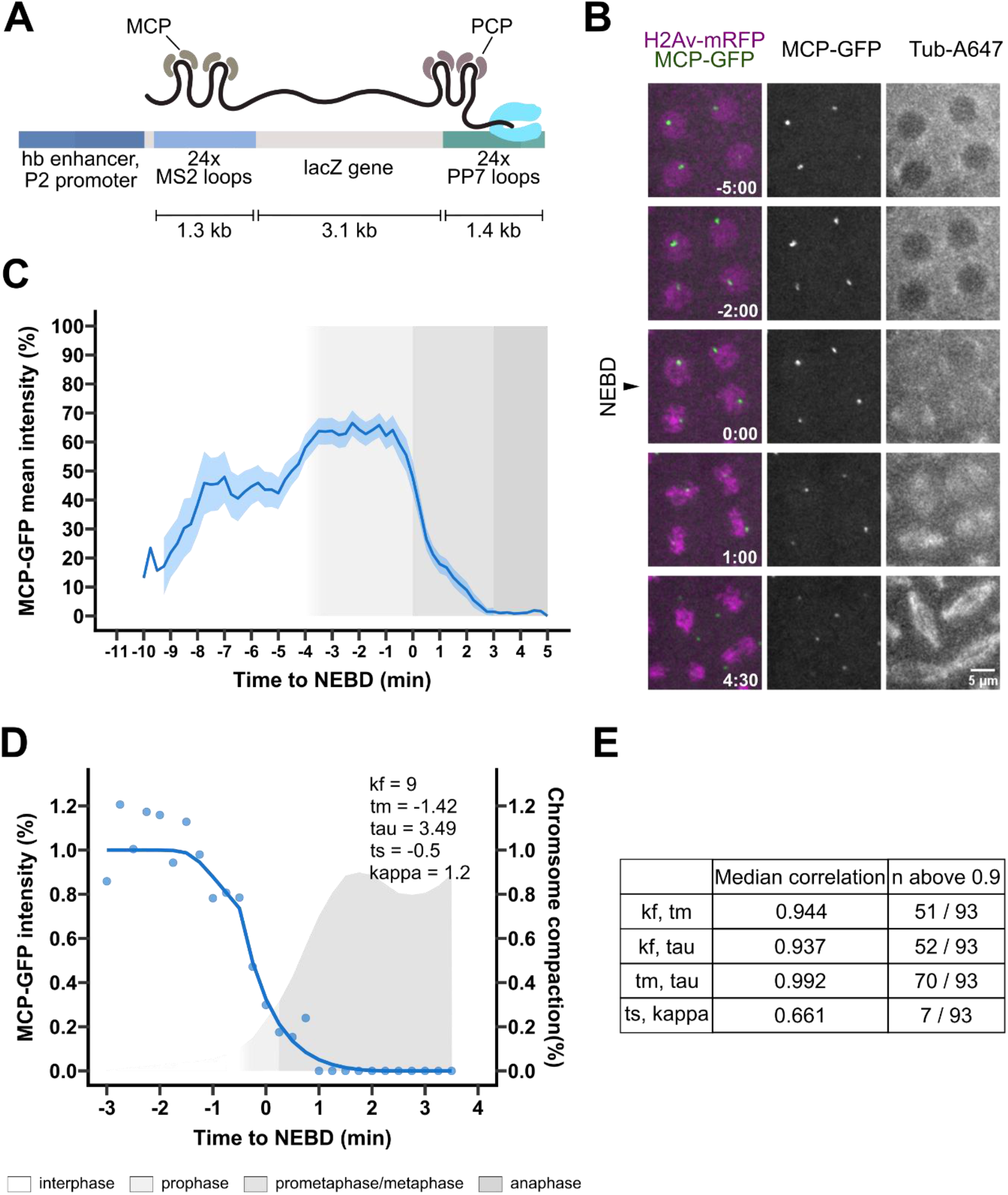
Transcriptional output and modelling relative to mitotic progression. A) Scheme of the transcriptional reporter used in this study. Adapted from (Fukaya et al., 2017). B) Nuclei of a Drosophila syncytial embryo at nuclear cycle 12 carrying the reporter in A and expressing His2Av-mRFP (magenta), MCP-GFP (green), and injected with tubulin-Alexa647. Tub-Alexa647 diffusion into the nuclear area (black arrowhead) is used to identify NEBD. Scalebar applies to all images. C) MCP-GFP normalized intensity over mitotic progression. Dark blue line mean, light blue area 95% Confidence Interval. Fluorescence intensity was normalized to the maximum value for each nucleus. All the tracks were aligned using NEBD as a reference time point. Grey boxes represent the mitotic stages as in the legend. N=93 nuclei from 11 embryos. D) Example of model fitting to an MCP-GFP track, r^2^ = 0.87. Blue dots, normalized intensity data, blue line, model fitting. Gray area, percentage of chromosome compaction over time, colored according to the mitotic stages as in panel C. E) Table showing the median correlation coefficient between pairs of parameters, and the number of nuclei with a correlation above 0.9. N= 93 nuclei from 11 embryos.

We next aimed to use the MCP curves to infer the kinetics and timings of the mechanisms underlying the repression of transcription during mitosis, focusing on the repression of new initiation events (Akoulitchev & Reinberg, 1998; Gebara et al., 1997; Long et al., 1998; Saad et al., 2025) and the postulated premature termination (Rothe et al., 1992; Shermoen & O’Farrell, 1991). To gain further insight into the kinetics of these events, we developed a mathematical model describing the fluorescent signal arising from the population of RNA polymerases present on the gene. At times preceding mitotic repression, approximately constant initiation maintains a baseline population of elongating polymerases, whose signal persists on chromatin for the duration of the elongation window, *τ* (tau), calculated as the ratio of the gene length over the elongation rate. When mitotic repression takes place, initiation of transcription decreases according to a sigmoidal function characterised by midpoint *t*_*m*_ and steepness *k*_*f*_, while polymerases already engaged on the gene continue elongating until they undergo canonical termination. The loss of the transcription signal is further accelerated by an active premature termination process, modelled as a delayed first-order reaction, occurring from time *t*_*s*_ with rate constant *κ* (kappa) (for details, see Materials and Methods and Figure S3).

We fitted the model to the decaying part of the MCP curves (Figure 1D). To limit parameter variation without strongly constraining the model, we imposed broad limits based on our experimental conditions and prior knowledge (Table S1). The model the fitted the data well, with a median r^2^ of 0.89 (IQR [0.83; 0.93]). However, analysis of the correlation index highlighted that *k*_*f*_, *t*_*m*_ and *τ* are highly interdependent (Figure 1E). This identifiability issue indicates that their individual kinetic contributions cannot be reliably resolved by fitting the transcriptional decay profiles alone, requiring independent experimental determination of these parameters.

### Transcription initiation is repressed gradually during mitosis

As the parameters controlling initiation decay cannot be estimated by model fitting, we wanted to determine experimentally when initiation is repressed and with which kinetics. The bulk MCP transcriptional signal is insufficient to understand how the single steps of the transcription cycle behave upon mitotic entry, as it results from the combination of multiple RNA polymerases at different transcriptional steps. To gain insight specifically into initiation kinetics, we adopted a FRAP (Fluorescence Recovery After Photobleaching)-based approach to estimate the presence/absence of new transcription events over time. The MS2-MCP binding is highly stable (Boireau et al., 2007; LeCuyer et al., 1996; Maiuri et al., 2011; Weil et al., 2010), and thus any recovery after FRAP can be used as a readout for the presence of new transcription initiation (Figure 2A). We confirmed that recovery after photobleaching is due solely to new transcription events and not to MCP turn-over, as recovery is not observed in the presence of triptolide (Figure S4), an inhibitor of transcription initiation (Bensaude, 2011).

**Figure 2.**
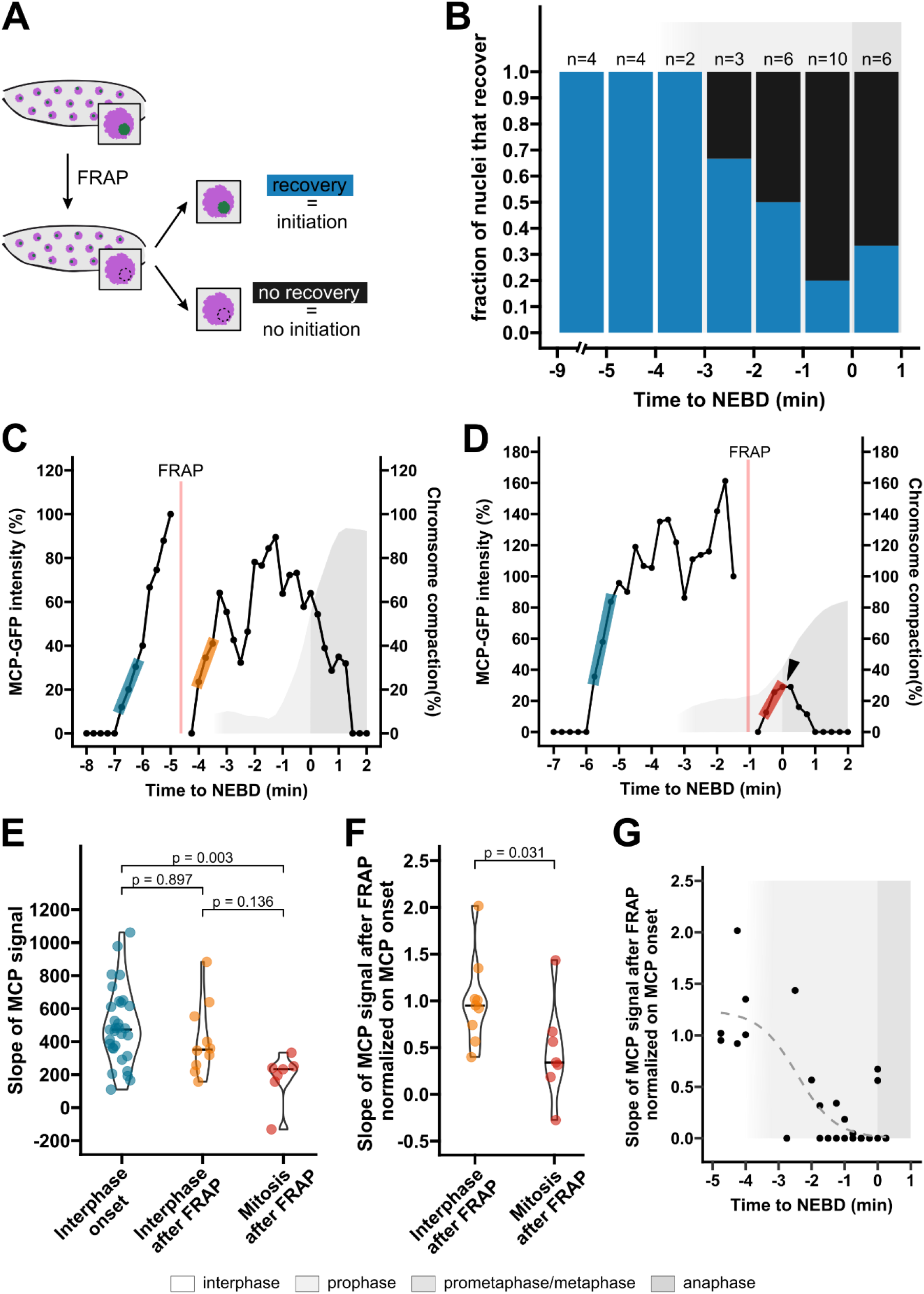
Repression of transcription initiation during mitosis is a gradual process completed around the time of NEBD. **A)** Experimental layout. MCP-GFP foci were photobleached, and their recovery was used to evaluate the presence of transcription initiation. **B)** Fraction of nuclei in which the MCP-GFP signal recovered after photobleaching, binned according to the time at which photobleaching was performed. Sample size as in figure. **C-D)** Example of MCP-GFP fluorescent intensityfrom nucleiphotobleached in interphase and mitosis, respectively. Blue, yellow and red segments highlight the slope of the curves at the onset of transcription and after photobleaching in interphase and in mitosis, respectively. Red vertical bar, FRAP. Grey areas represent the percentage of chromosome compaction coloured according to the mitotic stage as indicated. **E)** Slopes of the MCP signal calculated at the onset of transcription, after photobleaching in interphase, and after photobleaching in mitosis. Statistical analysis was performed using the nonparametric Kruskal–Wallis test (p = 0.0046) and post-hoc Dunn’s test for multiple comparisons with Bonferroni’s correction (adjusted p-values indicated in the panel). Only slopes with R squared > 0.65 were considered for analysis. **F)** Slopes after FRAP normalized on the slopes at the onset of transcription for each nucleus, grouped according to the cell cycle phase. Statistical analysis was performed using the nonparametric Wilcoxon test (p-value indicated in the panel). Only slopes with R squared > 0.65 were considered for analysis. Black lines indicated the median. **G)** Scatterplot of the normalized FRAP slopes plotted against the time at which FRAP was performed. The dashed grey line represents the fitting of the function describing initiation decay. For this analysis, nuclei where no recovery after FRAP was observed were also included (dots with a value of zero). Grey boxes, mitotic stages.

To evaluate how initiation of transcription is affected by cell cycle progression, we photobleached the MCP transcription signal at several time points (Figure 2A). We observed that FRAP during interphase was always followed by recovery of the transcription signal, consistent with the fact that transcription is active during this stage of the cell cycle (Figure 2B-C and S5A). When we performed photobleaching during prophase, we observed recovery after FRAP in a large fraction of embryos (Figure 2B, D and Figure S5B, C). FRAP after the time of NEBD resulted in recovery in a smaller fraction (0.3) of the nuclei (Figure 2B). These results highlight the presence of *de novo* initiation events even after cells enter mitosis, which are progressively repressed as mitosis advances.

To determine whether mitotic entry imposes any change in the rate at which RNA polymerases start transcribing, we estimated the rate of transcription initiation for each nucleus based on the slope of the 5’ MCP-GFP curves at the onset of transcription and after FRAP (Figure 2C-D, colored segments). We observed that in interphase, the recovery slopes are very similar to the ones at the onset of transcription (Figure 2E), with the slopes normalised to the onset of transcription assuming values around 1 (median 0.95, IQR [0.74; 1.02]) (Figure 2F), suggesting a constant initiation rate throughout interphase. In contrast, initiation rates estimated from nuclei in mitosis revealed a reduced rate (Figure 2E), with a median of 0.34, IQR [0.25; 0.61]) after intra-nuclear normalisation to the onset rate (Figure 2F). These findings reveal that entry into mitosis causes a decrease in the rate of transcription initiation.

To gain insight into the kinetics of this decrease, we fitted the normalised FRAP slopes with the sigmoidal function used in our model to describe progressive initiation repression. We also included the data points from embryos where no recovery was observed (initiation rate = 0) (Figure 2G). This analysis allowed us to experimentally determine the steepness *k*_*f*_ (1.71 ± 0.90 min^-1^, mean ± SEM, Standard Error of the Mean) and the midpoint *t*_*m*_ (-2.44 ± 0.48 min, mean ± SEM) of initiation repression (Figure 2G, grey dashed line). This means that the decay in initiation rate reaches its mid-point during prophase, and that the decay itself is shallow: with a of *k*_*f*_ =1.71, it takes approximately 3 min - almost the entire duration of prophase in this system - for the fluorescence to decay from 90% to 10% solely based on initiation inhibition.

Taken together, these results reveal that initiation is not abruptly inhibited as soon as cells enter mitosis, but the rate at which RNA polymerases start transcribing decreases as cells progress through mitosis, and *de novo* transcription is only prevented around NEBD.

### The elongation rate is unperturbed by mitotic progression

After establishing the kinetic parameters that govern changes in initiation rate relative to mitotic progression, we next aimed to experimentally estimate elongation rates and potential changes caused by mitotic entry. The interphase elongation rates were previously estimated for the *hb* reporter during nuclear cycle 13 and 14 as 2.4-2.6 kb/min (Fukaya et al., 2017). We followed a similar strategy to measure the elongation rate during nc12 (Figure S5). Analysis of the delay between the appearance of the 5’ MS2 and 3’ PP7 signals was used to determine the elongation rate in interphase 12, which was found to be slightly slower (1.94 kb/min, IQR [1.46; 2.12] kb/min) (Figure S6).

We next asked whether the elongation rate changes as cells approach mitosis. We followed the same FRAP-based strategy as to estimate initiation, but focusing on the initial recovery of 3’ transcription signal, as this is caused by RNA polymerases that are elongating and reaching the end of the gene, independently of new initiation events. We therefore used the slope of the 3’ fluorescence recovery as a proxy for the elongation rate. For this, we used a *hb* reporter in which the MS2 loops are located at the 3’ end (*hb-lacZ-MS2*) (Garcia et al., 2013) (Figure 3A, B), as PP7-PCP binding is not stably enough to exclude turn-over (Figure S4F-H). We estimated the slope of the 3’ MCP signal at the onset of transcription or after FRAP (Figure 3C, D). No difference in the slopes at the onset of transcription and after FRAP was detected (Figure 3E), even after intra-nuclear normalisation (Figure 3F). Therefore, the elongation rate in cycle 12 is ca. 2 kb/min and does not change during cell cycle progression. This further validates the assumption in our model that transcriptional silencing is driven mostly by changes in transcription initiation and premature transcription termination.

**Figure 3.**
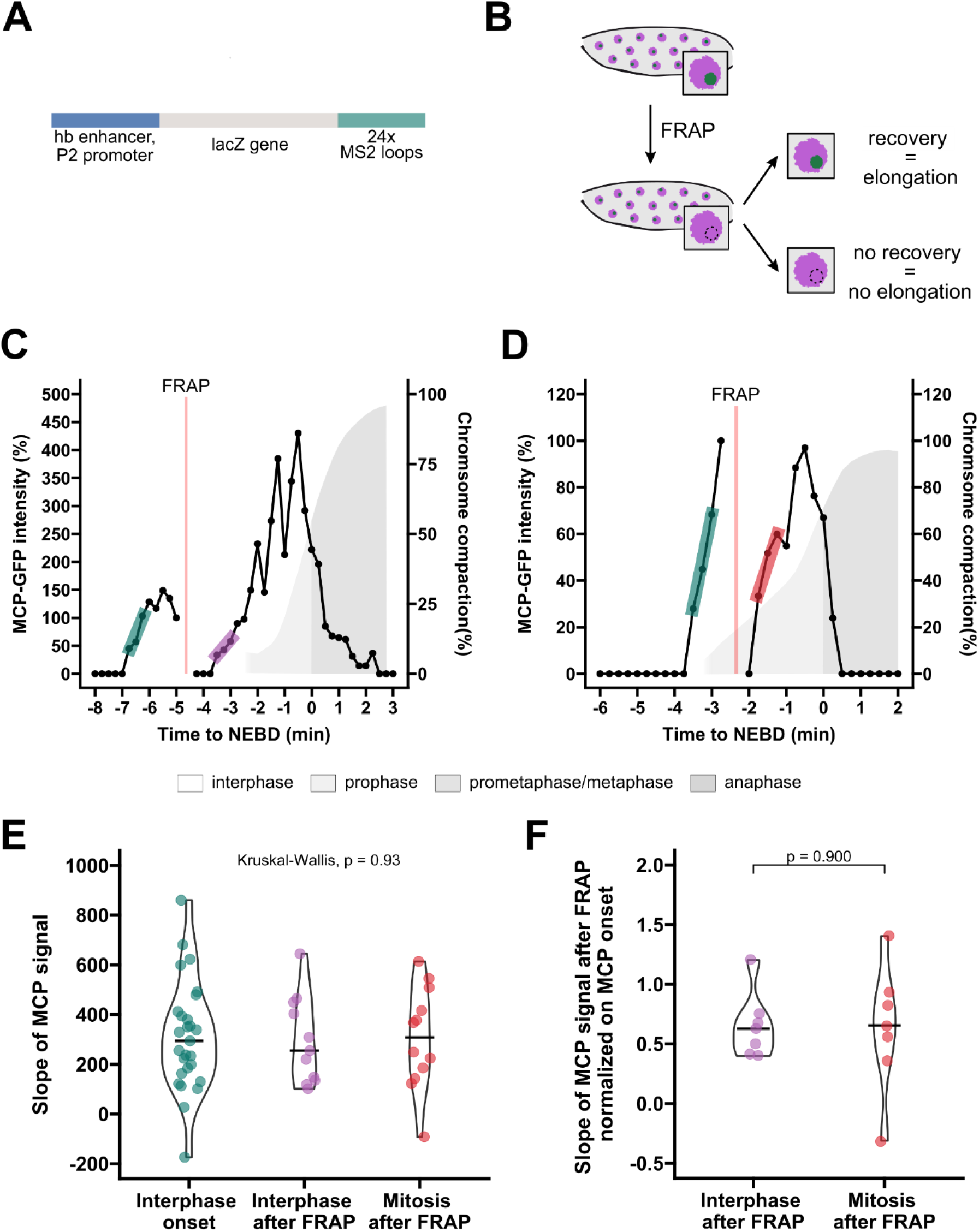
The elongation rate is not affected by cell cycle progression. **A)** Scheme of the reporter used, carrying 24xMS2 loops at the 3’. Adapted from (Garcia et al., 2013). B) Experimental layout. **C-D)** Example of 3’ MCP-GFP fluorescent intensity from nuclei photobleached in interphase and mitosis, respectively. Green, purple and red segmentshighlight the slope of the curves at the onset of transcription and after photobleaching in interphase and mitosis, respectively. Red vertical bar, FRAP. Grey areas represent the percentage of chromosome compaction coloured according to the mitotic stage as indicated. **E)** Slopes of the 3’ MCP signal calculated at the onset of transcription, after photobleaching in interphase, and after photobleaching in mitosis. Statistical analysis was performed using the nonparametric Kruskal–Wallis test (p-values indicated in the panel). **F)** Slopes after FRAP normalized to the slopes at the onset of transcription for each nucleus. Statistical analysis was performed using the nonparametric Wilcoxon test (p-value indicated in the panel). Only slopes with R squared > 0.65 were considered for analysis. Black lines indicate the median.

### Transcription is prematurely terminated during mitosis

The experiments in the previous sections allowed us to determine experimentally *t*_*m*_ and *k*_*f*_, the time and slope with which initiation is repressed during cell cycle progression. We also determined the elongation rate, from which we can derive *τ*, the time necessary to transcribe the entire gene. These are the three parameters that, by the initial fit of our model, were interdependent and not identifiable. Using the experimentally determined parameter windows (Table S2), we repeated the model fitting to estimate the time and strength of the premature termination mechanism,*t*_*s*_ and *K*. (Figure 4A). The fitting estimated premature termination to occur around 1 min after NEBD (*t*_*s*_, median, IQR [-0.04; 2] min) (Figure 4B) and with a strength *κ* around 9.2 min^-1^ (IQR [0.97; 9.2] min^-1^) (Figure 4C). Therefore, the premature termination mechanism acts in prometaphase and induces a sharp removal of the elongating polymerases from chromatin: in biological terms, a *κ* of 9.2 min^-1^ would induce a 90% reduction of transcription signal in just 15 sec. Notably, most of the nuclei (54/63) hit the upper bound imposed for *K*. This suggests that, for the majority of the nuclei, this parameter is at least as fast as 9.2 min^-1^, but the determination of the higher real value for this parameter was limited by our imaging resolution.

**Figure 4.**
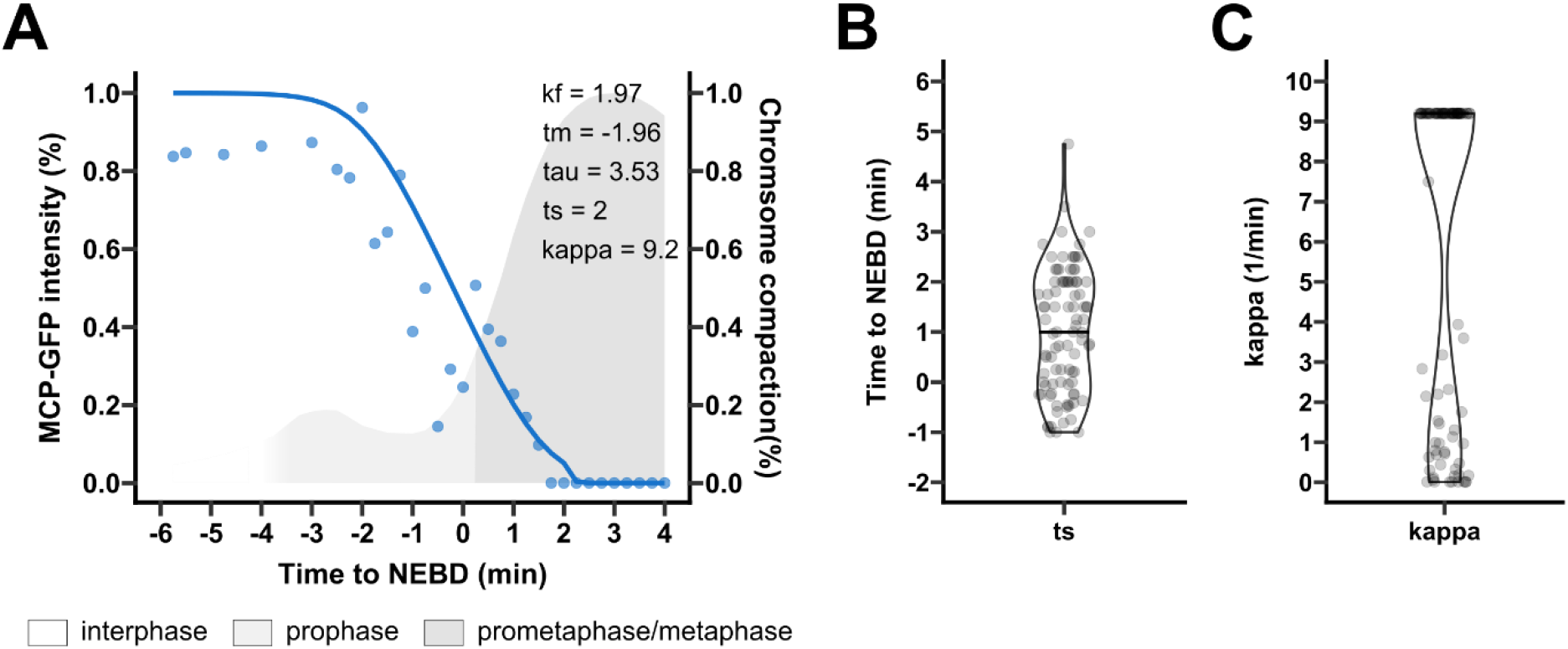
Premature termination removes acts on elongating complexes abruptly after NEBD. A) Example of model fitting to an MCP-GFP track. Kf, tm and tau were restricted as determined experimentally. Blue dots, normalized intensity data, blue line, model fitting. Gray area, percentage of chromosome compaction over time, colored according to the mitotic stages as in legend. B) Violin plot of the ts values estimated from the model fitting. Black line indicates the median. N= 93 nuclei from 11 embryos. C) Violin plot of the values of kappa estimated from the mathematical model. Black line indicates the median. N = 93 nuclei from 11 embryos.

Thus, our data, in combination with mathematical modelling, suggest that, while the initiation rate is slowly reduced during prophase, the complete repression of transcription during mitosis is achieved through a much stronger mechanism of premature termination that abruptly removes the elongating polymerases during prometaphase.

### Lds is responsible for premature termination of transcripts during mitosis

Having established the dynamics of mitotic transcriptional silencing, we next investigated its molecular basis. While the repression of initiation has long been attributed to Cdk1 (Cisek & Corden, 1989; Gebara et al., 1997; Leresche et al., 1996b; Long et al., 1998; Lüscher & Eisenman, 1992; Saad et al., 2025), the mechanisms behind premature termination are still unclear. The *Drosophila* helicase Lds/factor 2 has been shown to induce the release of nascent transcripts from RNA polymerases in *in vitro* studies (Liu et al., 1998; Xie & Price, 1997, 1996, 1998). Moreover, *in vivo*, the depletion of Lds induces chromatin retention of both the MCP signal (Carmo et al., 2023) and the Rpb3 subunit of RNA polymerase II during mitosis (Figure S7). In addition, our model predicts that the mechanism causing premature termination should become active around NEBD. Accordingly, quantitative analysis revealed that Lds binds chromatin at NEBD (median 0.0 min, IQR [-0.25; - 0.06] min) (Figure S8), in accordance with previous qualitative assessments (Carmo et al., 2023; Girdham & Glover, 1991). However, whether Lds works as a premature transcription termination factor and/or operates at other stages of the transcription cycle has not been established.

Retained transcription signals upon Lds depletion could be due to a delay in repressing transcription initiation. To test this possibility, we estimated the rate of initiation at mitotic entry (-3 to -0.5 min before NEBD), using FRAP of the 5’ MCP signal in Lds- and Luciferase-depleted embryos. This analysis revealed no change in the initiation rate upon Lds depletion (Figure S9). This finding excludes that Lds impacts the kinetics of transcription initiation and reinforces the notion that it may work at the premature termination stage. To directly evaluate this possibility, we performed imaging of the 5’ transcription signal from the *hb* reporter in Figure 1A in embryos where Lds had been depleted. We used the corresponding MCP curves to fit the mathematical model describing the decay of transcription as cells enter mitosis, restricting the parameters as estimated and reported in Table S3 (Figure 5 A-B). The onset of premature termination (*t*_*s*_) was significantly delayed in Lds RNAi embryos (Figure5 C). In addition, in Lds RNAi embryos, the median premature termination rate *K* has a value of 1.74 min^-1^ (IQR [0.01; 4.37] min^-1^), substantially lower than that estimate in control embryos (median 9.2 min^-1^, IQR [1.32; 9.2] min^-1^) (Figure 5 D). Moreover, the proportion of nuclei in which *K* is lower than 5 is statistically different between Lds- and Luciferase-depleted embryos (Lds 0.52, Luciferase 0.29, *X*^2^ = 12.33 with 1 degree of freedom, p-value=0.00022). These results indicate that depletion of Lds decreases the efficiency of premature termination, identifying Lds as a functional component of this active mechanism.

**Figure 5.**
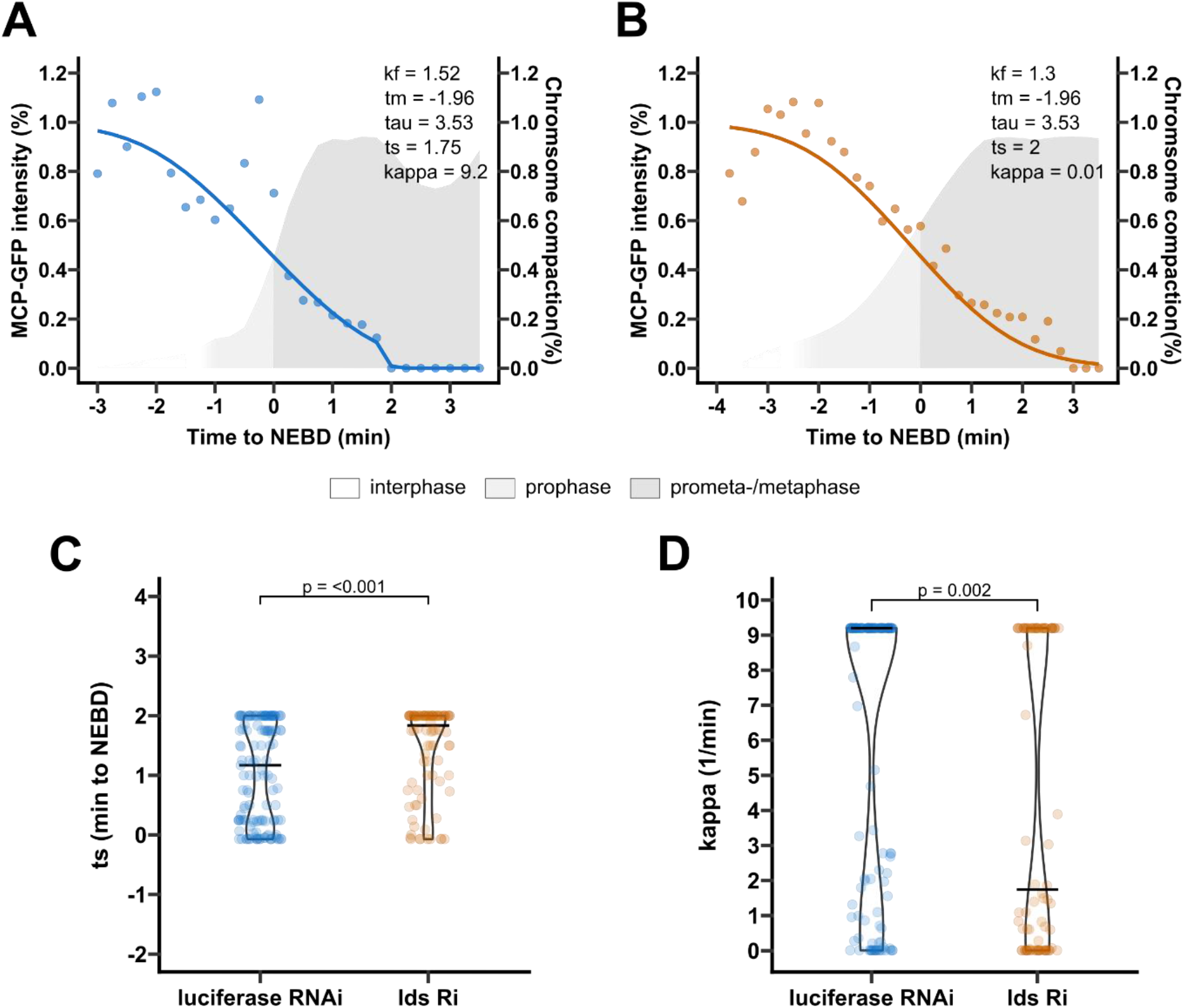
Lds is involved in premature termination of transcripts during mitosis. A) Example of model fitting to an MCP-GFP track from a Luciferase-depleted embryo (control). Kf, tm and tau were restricted as determined experimentally. Blue dots, normalized intensity data; blue line, model fitting. R^2^ = 0.86. Gray area, percentage of chromosome compaction over time, colored according to the mitotic stages as in legend. B) Example of model fitting to an MCP-GFP track from an Lds-depleted embryo. Kf, tm and tau were restricted as determined experimentally. Orange dots, normalized intensity data; orange line, model fitting. R^2^ = 0.94. Gray area, percentage of chromosome compaction over time, colored according to the mitotic stages as in legend. C) Violin plots of the ts values estimated from the model fitting for Luciferase (blue) and Lds RNAi embryos (orange). Black line indicates the median. Luciferase Ri, n = 129 nuclei from 10 embryos; Lds Ri, n = 88 nuclei from 10 embryos. Data were tested for normality using the Shapiro-Wilk test, and groups were compared using the non-parametric Wilcoxon test (p-value as in figure). D) Violin plots of the values of kappa estimated from the mathematical model for control Luciferase (blue) and Lds RNAi embryos (orange). Black line indicates the median. Luciferase Ri, n = 129 nuclei from 10 embryos, Lds Ri, n = 88 nuclei from 10 embryos. Data were tested for normality using the Shapiro-Wilk test and groups were compared using the non-parametric Wilcoxon test (p-value as in figure).

## Discussion

Although mitotic transcriptional silencing is thought to result from the coordinated action of multiple mechanisms (Contreras & Perea-Resa, 2024; Gottesfeld & Forbes, 1997), their relative contributions have remained poorly defined. Here, by complementing measurements of MS2/MCP transcriptional signal in dividing *Drosophila* embryos with FRAP, we quantitatively resolved the individual steps of the transcription cycle and determined how each is modulated during mitotic entry. We further used the experimentally determined transcriptional kinetics as parameters in a mathematical model describing the decay of transcription, from which we inferred further dynamics of the process of mitotic transcriptional silencing. We note that our model does not represent the individual molecular events underlying the silencing of transcription, *i*.*e*. which proteins interact and how; instead, it models the bulk effect that the mechanisms altogether have on the transcriptional output. Consequently, the inferred parameters should not be interpreted as direct measurements of individual molecular reactions, but as quantitative descriptions of the processes represented. For example, the parameter *k*_*f*_ describing initiation decay does not measure the absolute number of initiating RNA polymerases loaded on chromatin per unit of time, rather it describes that the number of initiating RNA polymerases experiences a shallow decay during the early stage of mitosis. Similarly, the rate of premature termination *K* does not measure the number of RNA polymerases removed from chromatin per unit of time, but it highlights that their removal happens at least as fast as our imaging resolution.

One of the key findings of this study is that transcription initiation for the *hb* reporter is not repressed abruptly upon mitotic entry but instead declines progressively over several minutes. This indicates that transcriptional silencing is regulated by a continuously changing input rather than by a single switch-like event. A plausible candidate is the gradual increase in mitotic Cdk1 activity, which has been shown to phosphorylate components of the transcriptional machinery in several models (Cisek & Corden, 1989; Gebara et al., 1997; Leresche et al., 1996b; Martínez-Balbás et al., 1995; Saad et al., 2025; Segil, 1996). Although the tight coupling between Cdk1 activation and mitotic progression makes this hypothesis difficult to test directly, the measured kinetics are consistent with initiation responding over a range of kinase activity rather than through an ultrasensitive threshold.

In contrast to transcription initiation, we found no evidence that RNA polymerase II elongation rate changes during mitotic entry. This suggests that ongoing elongation is largely insulated from the regulatory mechanisms that progressively silence transcription initiation, allowing engaged polymerases to complete transcription at a constant rate during the early stages of mitosis. Interestingly, this means that the speed of the RNA polymerases is unaffected by the changes in the chromatin that happen during prophase.

Our modelling further supports that transcriptional silencing requires active removal of elongating RNA polymerases during prometaphase, as already proposed in (Rothe et al., 1992; Shermoen & O’Farrell, 1991), and identifies Lds as an important contributor to this process. We did not find any evidence of Lds affecting initiation dynamics, though a cross-talk between initiation repression and premature termination may exist, possibly through Cdk1 activity. Importantly, delayed transcript removal upon Lds depletion has also been observed with additional reporter constructs (Carmo et al., 2023), supporting a broader role for Lds. This suggests that the regulatory dynamics described here may not be unique to the reporter used in this study, though we cannot exclude that genomic context or locus-specific properties may modulate the precise kinetics of this process. In addition, depletion of the Lds homolog TTF2 in human cells leads to the accumulation of nascent transcripts throughout mitotic chromatin (Jiang et al., 2004; Tovini et al., 2026). These observations support a conserved role for Lds/TTF2 in mitotic transcript clearance and suggest that the regulatory mechanisms described here are neither unique to our reporter nor a specialisation of rapidly dividing embryonic cells. Nevertheless, their precise kinetics may vary with developmental, cellular, or genomic context.

Indeed, although mitotic transcriptional repression is broadly conserved across metazoans (Davidson, 1964; Taylor, 1960), the relative contribution and dynamics of the processes that enforce silencing may differ between cell types and species. Consistent with this idea, TTF2 depletion produces markedly different patterns of mitotic transcript retention in HeLa and H9 cells, with substantially greater persistence of nascent mRNA transcripts through metaphase in HeLa cells (Pedro et al., 2026). This suggests that the kinetics and relative contribution of TTF2-dependent clearance can vary between cellular contexts.

Additional regulatory processes beyond the ones described here may operate in specific cellular or transcriptional contexts. In Hela cells, for example, enhanced promoter escape contributes to mitotic genome silencing by releasing RNA polymerases stalled at promoters (Akçaöz-Alasar et al., 2026; Liang et al., 2015). This mechanism is unlikely to take place in the *Drosophila* embryo, as most early zygotic genes, including the *hunchback* promoter used in our work, do not exhibit promoter-proximal pausing during interphase at the blastoderm stage (Chen et al., 2013). Thus, while the phenomenon of mitotic transcriptional silencing is broadly conserved, its precise implementation is likely shaped by the transcriptional and cellular context.

The existence of multiple, conserved pathways that repress transcription during mitosis underscores the biological relevance of this process (Davidson, 1964; Feinendegen et al., 1960; Fink & Turnock, 1977; Shermoen & O’Farrell, 1991; Taylor, 1960). Failure to silence transcription can interfere with chromosome organisation and segregation (Carmo et al., 2023; Liang et al., 2015; Perea-Resa et al., 2020; Tovini et al., 2026), while mitotic transcriptional regulation has also been implicated in the control of cell identity and developmental gene expression (Contreras & Perea-Resa, 2024; Halley-Stott et al., 2014). By resolving the distinct dynamics underlying transcriptional shutdown, our work lays the ground for selectively perturbing these processes and dissecting their contributions to chromosome segregation, cell fate and development.

## Materials and Methods

### Fly stocks

*Drosophila melanogaster* populations were kept at 18ºC or 25ºC in vials of Vienna fly food. The genotypes used are reported in Table S4.

### Embryo preparation and microinjections

Syncytial embryos were collected every 1:30-2 hr in apple juice agar plates with yeast paste. Embryos were prepared for imaging as previously described (Carmo et al., 2019). Briefly, embryos were incubated for 2 min in 50% bleach, rinsed with water, and aligned on a piece of agar. Embryos were subsequently attached to a 24 × 40 mm coverslip coated with a thin layer of heptane glue, dried for 13 min, and covered with Halocarbon oil 700. Microinjections were performed at the anterior pole of the embryos using an Eppendorf FemtoJet Microinjector and Eppendorf Femtotip sterile injection capillaries (0.5 μm inner and 1.0 μm outer diameter). Compounds were injected at the following concentrations: 1.5 µg/µl porcine brain tubulin-HiLyte647 in 1M BRB80 at pH 7.8; 500 µM triptolide dissolved in DMSO; 10% DMSO in distilled water; 0.5 µg/µl WGA-Alexa647 in 1X PBS. Tubulin was injected in all embryos expressing MCP-GFP before imaging. Triptolide and DMSO were injected in single embryos after confirming expression of the MCP-GFP signal and were co-injected with WGA-Alexa647 to monitor the point of injection. Embryos at nc12 were identified based on nuclear density.

### Microscopy

Experiments requiring FRAP and/or microinjections were performed in a Yokogawa CSU-X Spinning Disk confocal, mounted on a Leica DMi8 microscope, with a 63 × 1.3NA glycerine immersion objective, using the 561, 488 and 637 laser lines and an Andor iXon Ultra EMCCD 1024 × 1024 camera. The system was controlled with Metamorph software (Molecular Devices). For every channel, 6 z-stacks were acquired every 0.7 µm, at intervals of 15 sec. To estimate the elongation rate in nc12, imaging of MCP-GFP and PCP-mCherry was performed on a Zeiss LSM 900, with a 40x water immersion objective. For each channel, 14 stacks were acquired every 0.7 µm, at time intervals of 8.97 sec. The laser powers were previously calibrated using a *hb-MS2-PP7-lacZ* reporter to ensure that the MCP and PCP signals would arise at the same time and have similar dynamics.

### Fluorescence Recovery After Photobleaching (FRAP)

FRAP experiments were performed using an Andor Mosaic system. Embryos were imaged as above, starting at metaphase or early interphase. When the MCP-GFP signal was visible in several nuclei, the acquisition was stopped, and a single MCP-GFP dot was selected with circular ROI of 2.19 µm diameter. Photobleaching of the ROI was performed by applying 1 pulse of a 470 nm laser at 60% power for 200 ms. Only one MCP dot per embryo was photobleached.

### Quantitative imaging analysis

All images acquired were max-projected using Fiji. Nuclei of interest were cropped, and the MCP-GFP dots were followed over time using TrackMate (Ershov et al., 2022). Signal intensity was measured until the foci were displaced from chromatin. For every time point, a background measurement was performed using a circular ROI placed on the nucleus but not overlapping with the MCP signal. For time points where no MCP foci were detected on chromatin, the background values were used.

### Characterization of cell cycle progression in *Drosophila* embryos

To characterize mitotic progression in *Drosophila* embryos at nc12, we performed live imaging of embryos expressing His-RFP, PCNA-GFP and injected with tubulin-Alexa647 (Figure S1A). End of S-phase was defined by the time of PCNA-GFP foci disappearance. As at this time most nuclei reached 10% chromosome compaction (see below), this percentage was used to identify mitotic entry. NEBD was identified by the diffusion of tubulin in the nuclear area and anaphase onset by the poleward movement of chromosomes. Estimation of chromosome compaction over time was obtained by first creating a mask on the histone channel with low intensity threshold to cover the entire nuclear area at all time points. Then, a second mask was created with a higher intensity threshold to cover just the bright pixels of compacted chromatin. The resulting data on percentage of compacted chromatin were smoothed using the loess method to create a continuous distribution and calculate the time at which 10% compaction was reached.

### Calculation of the MCP signal slopes

For each photobleached nucleus, we estimated the slope of the MCP curves by calculating the linear regression coefficient at the onset of transcription and after FRAP. The linear regression was performed over the first 3 data points. Considering such a short interval was necessary to rule out any transcript leaving chromatin, which would otherwise cause an underestimation of the rate. Any regression with an R-squared lower than 0.65 was not considered.

### Calculation of the elongation rate

The elongation rate during interphase was calculated as in (Fukaya et al., 2017). Briefly, embryos carrying His-eBFP, PCP-mCherry, MCP-GFP, and a *hunchback* reporter carrying interspersed 12XMS2 and 12xPP7 at the 5’ were imaged to determine the laser powers at which the MCP and PP7 signals arise at the same time and behave with similar dynamics. The same imaging conditions were used to image embryos carrying the *hb-MS2-lacZ-PP7* reporter. These embryos were imaged from mitosis 11 to interphase 14. Nuclei expressing both MCP and PCP signals were selected and individually processed as described above. The MCP-GFP and PCP-mCherry tracks obtained were smoothed using the LOESS (Locally Estimated Scatterplot Smoothing) method. The time points at which the two tracks reached 30% of the maximum intensity were determined and used to calculate the lag time. The elongation rate was calculated by dividing the length of the *MS2-lacZ-PP7* reporter (4527 bp) by the lag time.

### Statistical analysis

Data were analyzed with R Statistical Software (R Core Team, 2023), the *readxl* (v1.4.5; Wickham H., Bryan J., 2025), *writexl* (v.1.5.3; Ooms J., 2025), *dplyr* (v1.3.1; Wickham H., François R., Henry L., Müller K., Vaughan D., 2023), *minpack*.*lm* (v1.2-4; Elzhov T.V., Mullen K.M., Spiess A., Bolker B., 2023), *ggplot2* (Wickham H., 2016) and *ggprism* (v1.0.7; Dawson C., 2025) packages. All datasets were tested for normality using Shapiro-Wilk normality test and alpha=0.05. Parametric or nonparametric tests were performed considering the normality of the samples. When performing multiple comparisons, the alpha value was adjusted accordingly. Details for each test can be found in the respective figure legends.

### Mathematical modelling

The mitotic decay in initiation events was described as a sigmoidal function, which decays over time with a rate *k*_*f*_ and reaches half of its decay at time *t*_*m*_:

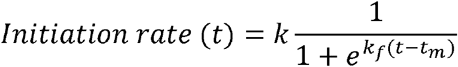

With *k* scaling factor. The elongating RNA polymerases will leave chromatin after an elongation window *τ*(tau), given by the ratio of the gene length and the elongation rate. Therefore, the fluorescence signal over time *F*(*t*) is given by the RNA polymerases that start transcribing at a decreasing rate and the ones that leave chromatin due to canonical termination. Mathematically, this corresponds to the integral of the initiation decay function over the elongation window:

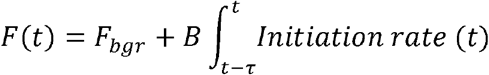

Where *F*_*bgr*_ is the fluorescence background and *B* a scaling factor. The integral can be solved analytically by defining:

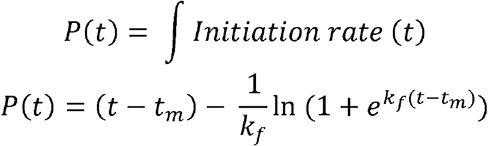

The fluorescent signal becomes:

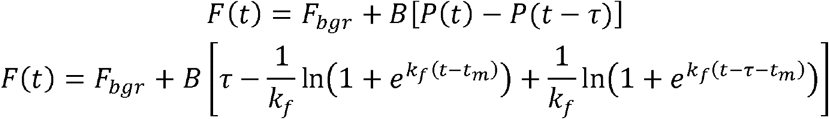

Before *t*_*m*_, this equation has a plateau at value *B* · *τ*. This makes the two parameters not identifiable, as changes in one would be compensated by changes in the other. To solve this issue, we normalized by *τ*. In addition, as the MCP tracks were background-subtracted and normalized to the maximum, we assumed *F_bgr_* = 0 and *B*=1. Thus, the equation becomes:

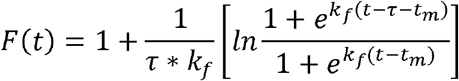

This equation describes the decay in fluorescent signal as given solely by the gradual reduction of RNA polymerases that start transcribing and their removal from chromatin due to canonical termination. In the presence of an additional premature termination mechanism, the RNA polymerases will be suddenly removed from chromatin at a time *t*_*s*_ and with strength *κ* (kappa). This modifies the previous equation as follows:

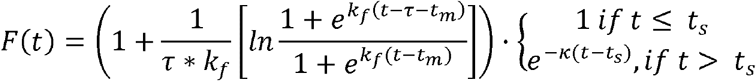

This equation was fit to the data in R, and variables were constrained as described in Tables S1-S3. As the equation describes only the decay of transcription, the MCP tracks were cut 5 time points before the maximum. A number of zeros equal to half the length of the track was added at the end to ensure an equally good fitting across tracks independently of their length. The fluorescence was expressed relative to the time of NEBD. Each nucleus was fitted independently using the Levenberg– Marquardt algorithm (nls.lm, R package *minpack*.*lm*), with convergence criteria maxiter = 500, ftol = ptol = 1e-8. The best starting values were determined both with a grid search and manual testing of specific combinations. To minimise the chance of converging to a local minimum, each fitting was restarted 50 times from different starting values, and the fit with the lowest SSE was considered for downstream analysis.

For each nucleus independently, the parameter correlation matrix was computed from the Jacobian matrix *J* of the model residuals derived numerically at the best-fit parameter values (using the *numDeriv* package in R). The parameter covariance matrix was estimated as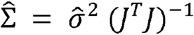, where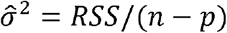 is the residual variance (*n* number of time points, *p* number of parameters, *RSS* Residual Sum of Squares). From the covariance matrix, the linkage between the parameters was standardized into a correlation matrix using 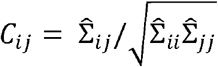. For each parameter pair, we reported in the table the median absolute correlation and the percentage of nuclei exceeding a threshold of |0.9| across all nuclei.

## Declaration of generative AI and AI-assisted technologies in the manuscript preparation process

During the preparation of this work, the authors used ChatGPT (OpenAI) in order to improve the readability and language of the manuscript, and Claude (Anthropic) as an assistance tool for mathematical modelling, including coding and troubleshooting. The authors reviewed and edited the output as needed and take full responsibility for the content of the published article.

## Authors Contributions

P.G.: Conceptualisation, Methodology, Investigation, Visualisation, Writing (original graft); A.T.: Resources, Investigation, Writing (review & editing). T.P.: Methodology, Writing (review & editing) R.A.O.: Conceptualisation, Methodology, Writing (original draft), Supervision, Funding acquisition.

## Acknowledgements

R.A.O. dedicates this work to the memory of Béla Nóvak, who introduced her to quantitative biology and mathematical modelling—an influence that shaped this study. We would like to thank present and past members of the Oliveira lab for fruitful discussions and feedback. We also thank Bloomington Drosophila Stock Center (BDSC), Patrick O’Farrell (University of California), Michael Levine (Princeton University) and João Raimundo (Católica Biomedical Research Centre) for sharing fly stocks. We acknowledge the support of the Bioimaging Platform of the Gulbenkian Institute for Molecular Medicine, funded by PPBI-POCI-01-0145-FEDER-022122, and of the GIMM Fly Facility. This work was funded by the European Research Council (ERC) under the European Union’s Horizon 2020 research and innovation programme (grant agreement No 101002391) awarded to R.A.O., and FCT - Fundação para a Ciência e Tecnologia, I.P. by project reference UI/BD/154413/2022 and DOI identifier https://doi.org/10.54499/UI/BD/154413/2022 supporting P.G.

