## Supplementary material for "Distinct regulation across the transcription cycle drives mitotic transcriptional silencing": Figure S1

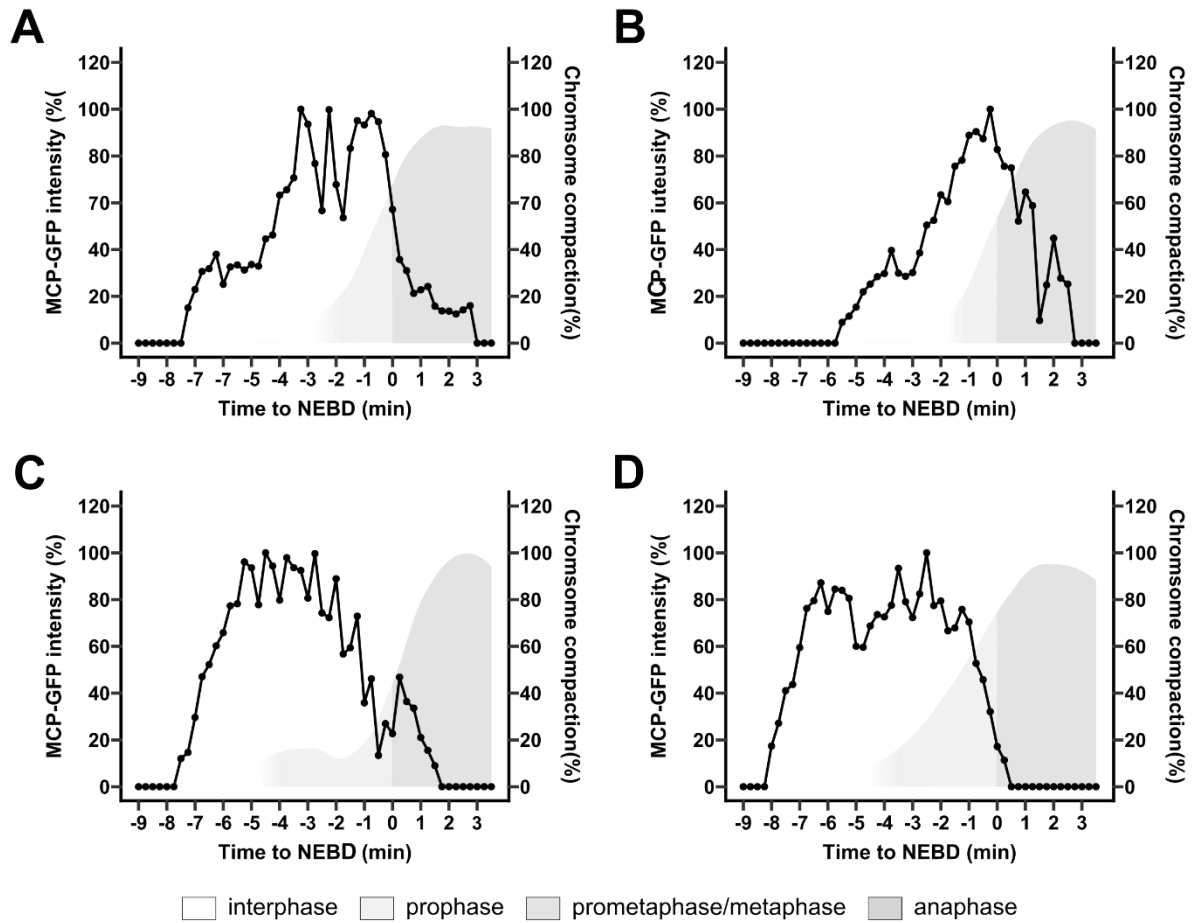

**Figure S1. Single MCP-GFP transcription tracks along the cell cycle.** A-D) Example of MCP-GFP mean intensity over time in single nuclei. Grey boxes indicate the progression through the mitotic stages as determined in Supplementary Fig2.
