## Supplementary material for "Distinct regulation across the transcription cycle drives mitotic transcriptional silencing": Figure S2

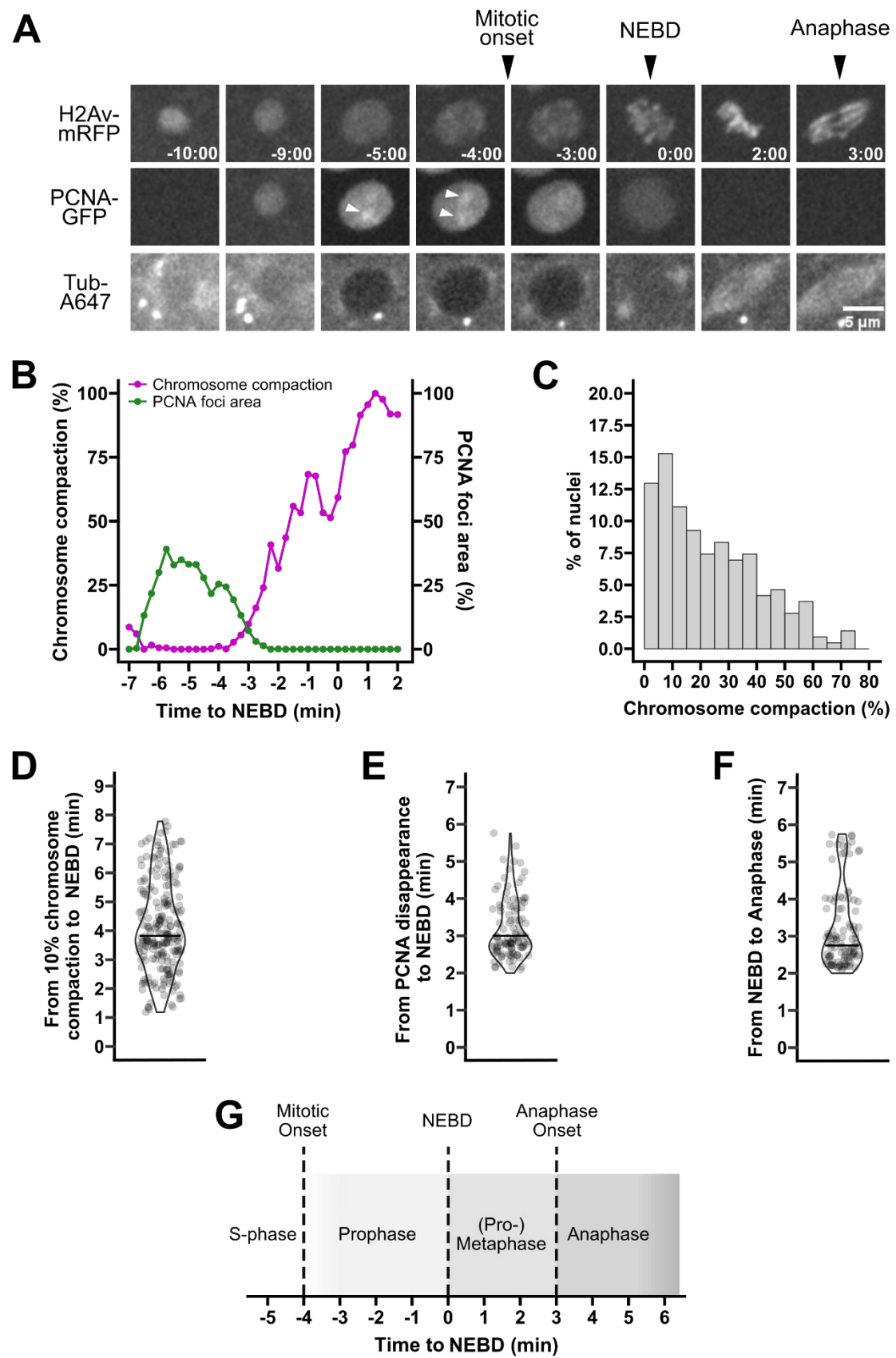

**Figure S2. Characterization of mitotic progression in *Drosophila* embryos at nuclear cycle 12.** A) Nucleus expressing His2Av-mRFP, PCNA-GFP and injected with tubulin-Alexa647. PCNA forms nuclear patches (white arrows) that disappear as nuclei enter mitosis. NEBD can be

identified as the moment in which tubulin invades the nuclear area. Anaphase can be assessed by the separation of chromosomes. B) Example of chromosome compaction over time against the area occupied by the PCNA patches for one nucleus. C) Histogram of the percentage of nuclei showing specific intervals of chromosome compaction at the time of PCNA disappearance. N = 216 from 5 different embryos. D) Time from 10% chromosome compaction to NEBD evaluated by tubulin staining. Black bar represents the median. Median 3.82 min, Inter Quartile Range (IQR) [3.13; 5.17]. N = 216 from 5 different embryos. E) Time from disappearance of the PCNA patches to NEBD evaluated by tubulin staining. Black bar represents the median. Median 3 min, IQR [2.75; 3.75]. N = 105 nuclei from 7 different embryos. F) Time from NEBD evaluated by tubulin staining to anaphase evaluated from the separation of chromosomes. Black bar represents the median. Median 2.75 min, IQR [2.5; 3.75]. N = 105 nuclei from 7 different embryos. G) Reference time scheme for mitotic progression in *Drosophila* embryos at nc12. Data was tested for normality using the Shapiro-Wilk normality test.
