## Supplementary material for "Distinct regulation across the transcription cycle drives mitotic transcriptional silencing": Figure S3

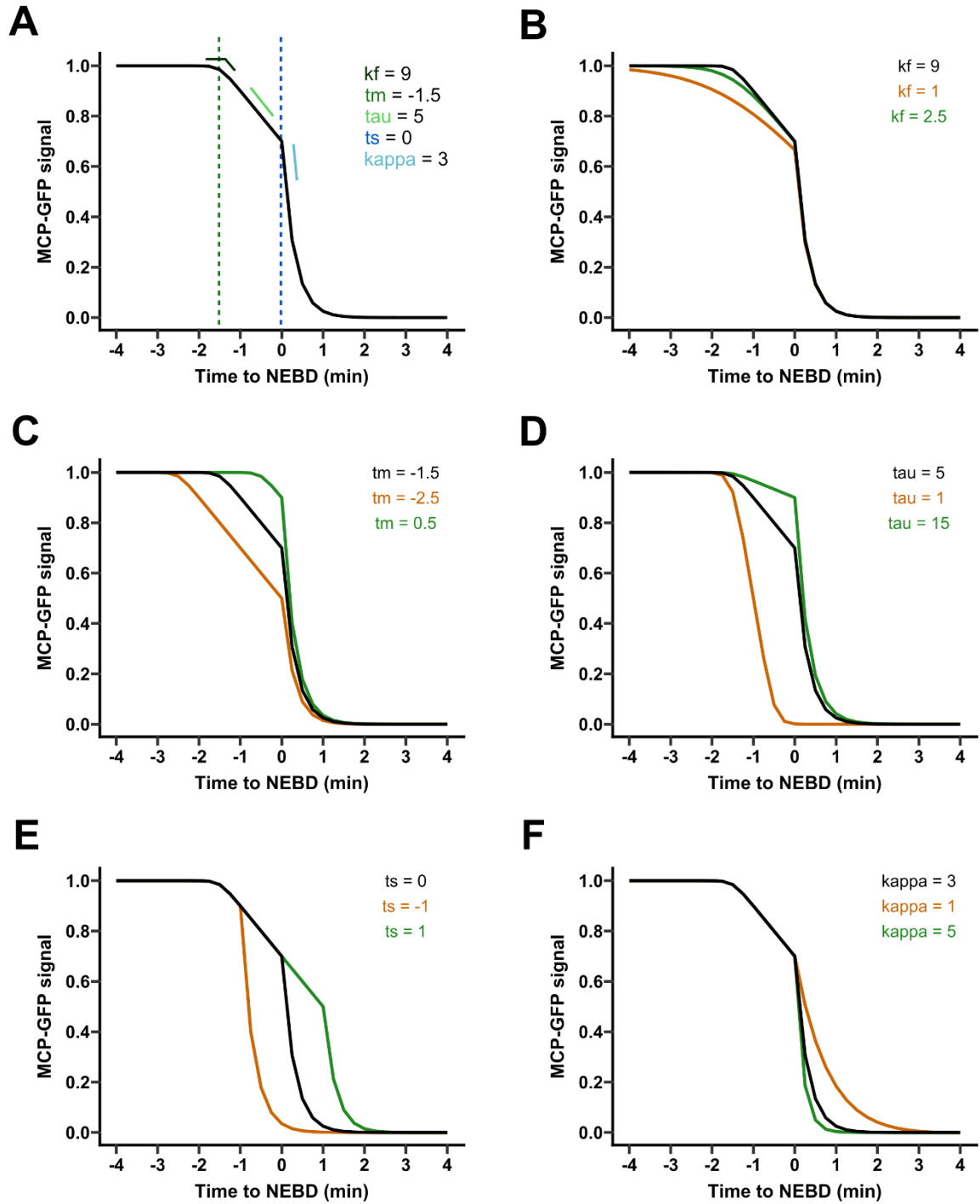

**Figure S3. Mathematical model that describes the decay of transcription as nuclei enter mitosis.** A) Example of a curve obtained with the parameters shown in figure. Colored lines highlight the contribution of each of the parameters. B-F) Example of curves where one parameter at a time was changed to highlight the different contributions of each parameter to the shape of the curve.
