## Supplementary material for "Distinct regulation across the transcription cycle drives mitotic transcriptional silencing": Figure S4

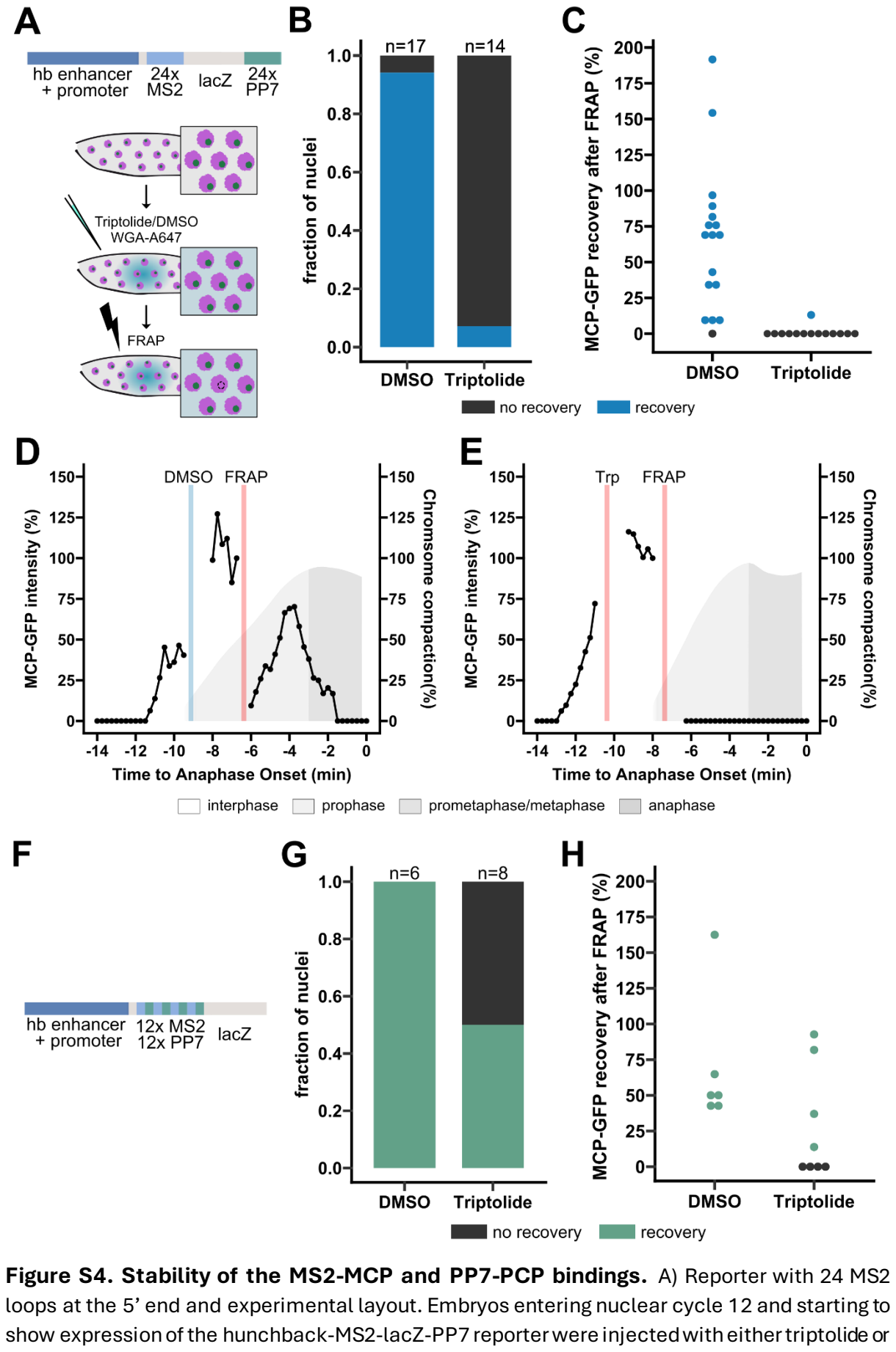

Figure S4. Stability of the MS2-MCP and PP7-PCP bindings. A) Reporter with 24 MS2 loops at the 5’ end and experimental layout. Embryos entering nuclear cycle 12 and starting to show expression of the hunchback-MS2-lacZ-PP7 reporter were injected with either triptolide or

DMSO. WGA-Alexa647 was co-injected as a marker to localize the injection spot. For a nucleus in the injected area, the MCP-GFP focus was photobleached. Injection and photobleaching were performed as early as possible to give enough time for the MCP foci to recover before mitosis. B) Evaluation of the fraction of nuclei in which the MCP-GFP signal recovered after photobleaching, binned according to the treatment. C) Percentage of recovery of the MCP-GFP signal after photobleaching, binned according to the treatment. D, E) Examples of MCP-GFP intensities in nuclei that were photobleached after injection of DMSO and triptolide, respectively. F) Reporter carrying 12 PP7 loops at the 5' end. G) Evaluation of the fraction of nuclei in which the PCP-GFP signal recovered after photobleaching, binned according to the treatment. H) Percentage of recovery of the PCP-GFP signal after photobleaching, binned according to the treatment. In contrast to what was observed for the MS2-MCP binding, the PCP signal recovers after FRAP in the presence of triptolide for several nuclei, highlighting the instability of the PP7-PCP binding.
