## Supplementary material for "Distinct regulation across the transcription cycle drives mitotic transcriptional silencing": Figure S5

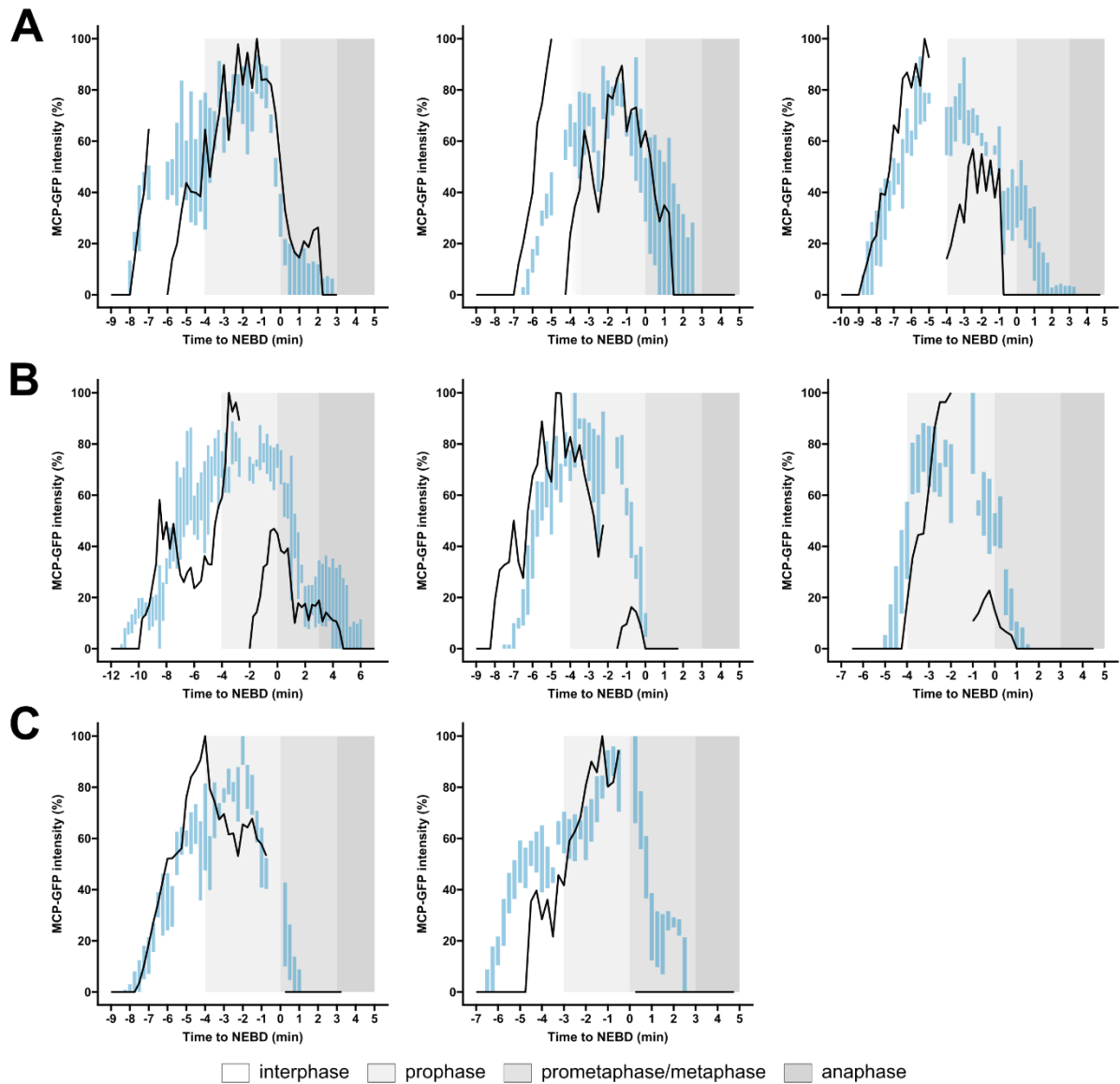

**Figure S5. Additional examples of 5' MCP FRAP.** Examples of FRAP of the 5' transcription signal in interphase (A), early prophase (B) and late prophase (C). Black line represents the MCP intensity from the photobleached MCP spot, normalized to the maximum. Blue bars represent the interquartile distribution of the MCP fluorescence from 4 neighbor nuclei that were not photobleached, normalized to the maximum. Grey areas represent the mitotic stages as in the legend.
