## Supplementary material for "Distinct regulation across the transcription cycle drives mitotic transcriptional silencing": Figure S6

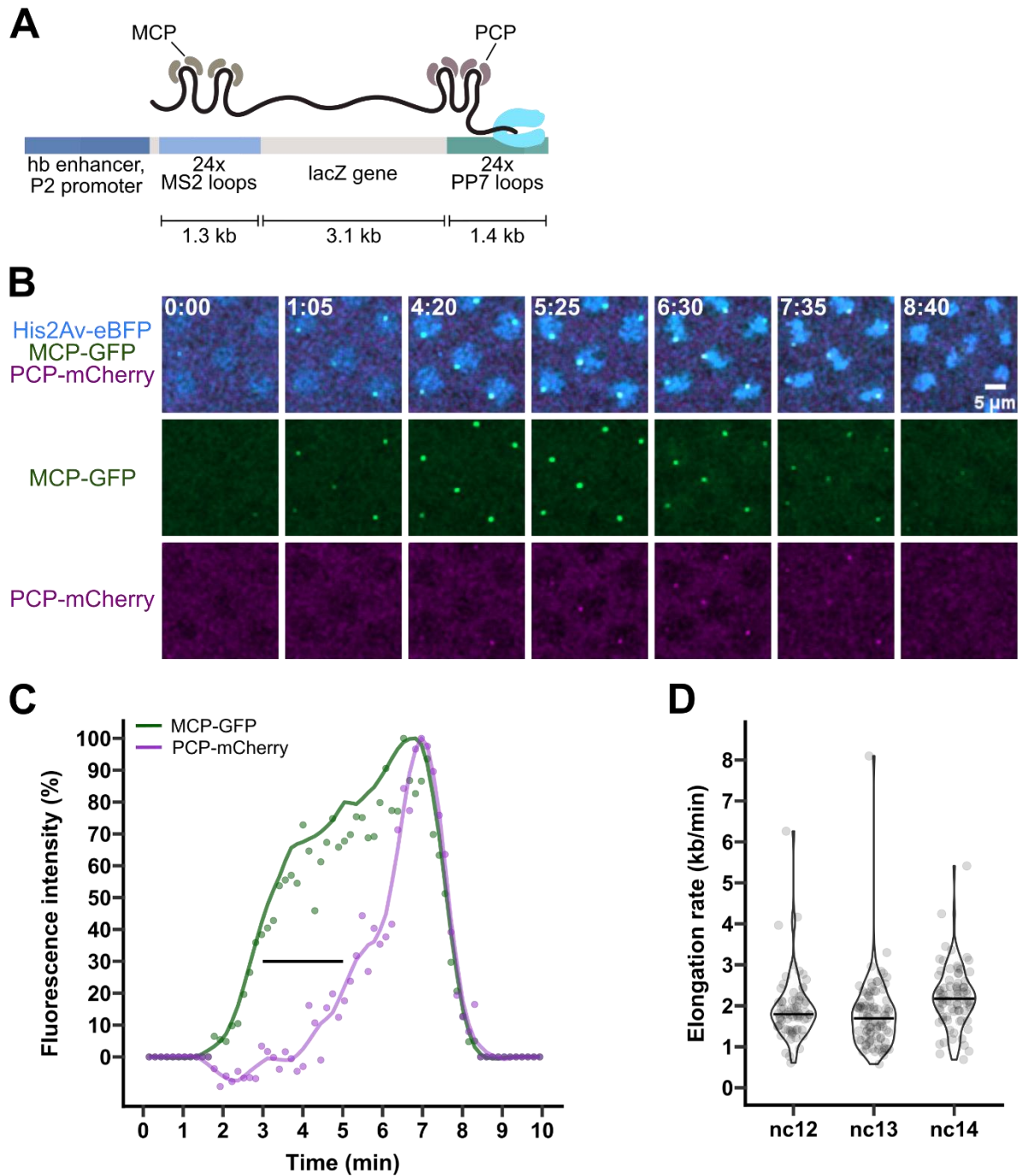

**Figure S6. Elongation rate during interphase.** A) Scheme of the reporter used for this experiment. B) Example of nc12 nuclei expressing His2Av-eBFP, MCP-GFP and PCP-mCherry. Scale bar applies to all images. C) Example of MCP-GFP (green dots) and PCP-mCherry (magenta dots) intensity over time. The data were smoothed with the loess method (green and magenta lines), and the time at which the two tracks reached 30% of their fluorescence intensity was determined. The delay between these time points (black horizontal line) was used to calculate the elongation rate. D) Elongation rate per nucleus in embryos at nc12, nc13 and nc14. Nc12, n = 90 nuclei from 4 different embryos. Nc13, n = 103 nuclei from 5 embryos. Nc14, n = 81 nuclei from 4 different embryos.
