## Supplementary material for "Distinct regulation across the transcription cycle drives mitotic transcriptional silencing": Figure S7

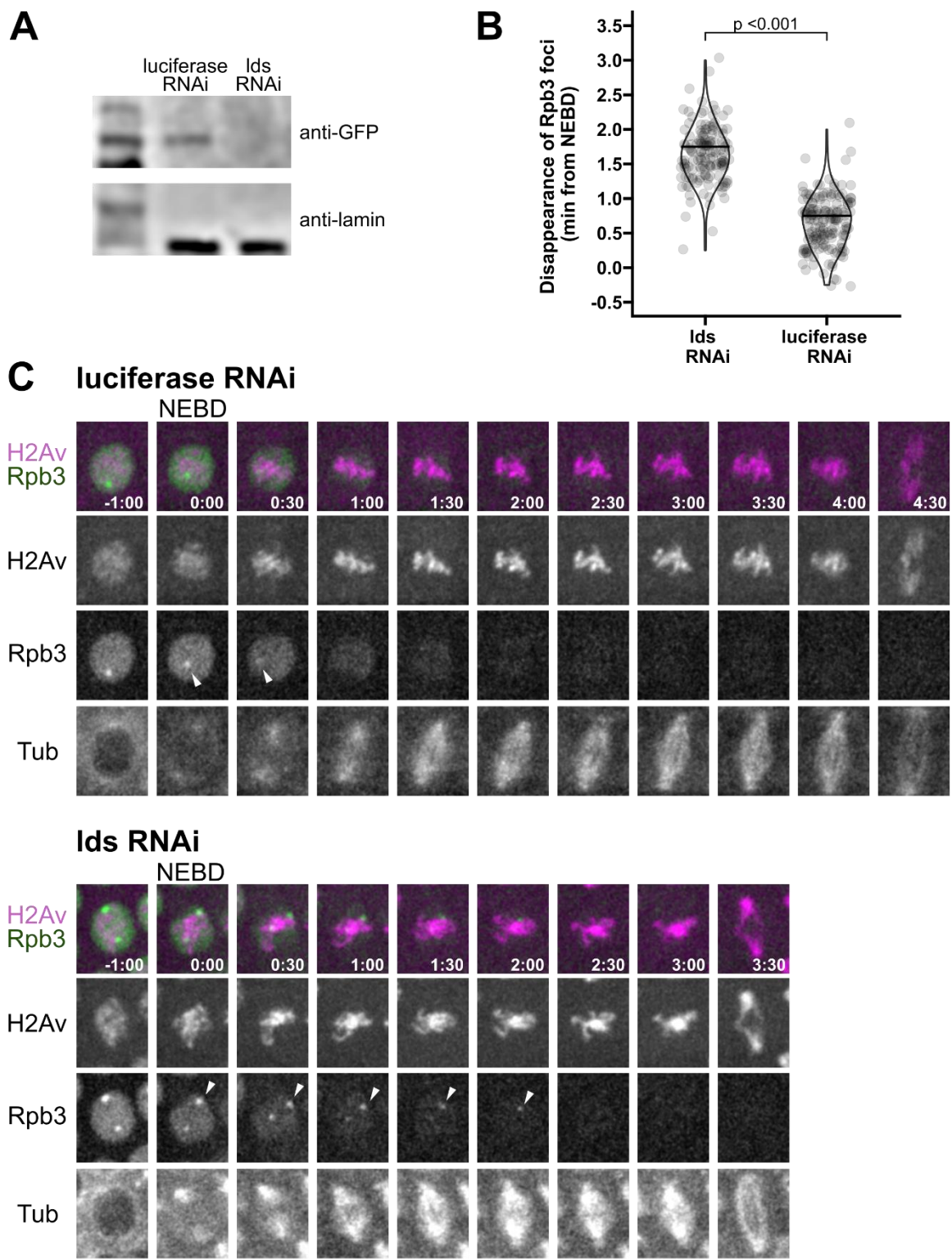

**Figure S7. Depletion of Lds by RNAi is complete in embryos and causes retention of Rpb3 on mitotic chromatin.** A) Western Blot showing depletion of Lds in luciferase RNAi and Lds RNAi expressing embryos. Lds-GFP was detected with an anti-GFP antibody. Loading control, lamin. B) Quantifications of the time that Rpb3 foci are retained in Lds

and luciferase RNAi embryos. Statistical analysis was performed using the nonparametric Wilcoxon test (p-value $8.33\text{e-}44$). Sample size 141 nuclei from 7 lds RNAi embryos, 158 nuclei from 8 luciferase RNAi embryos. C) Examples of control or lds-depleted embryos expressing H2Av-mRFP, EGFP-Rpb3 and injected with tubulin-Alexa647. Time is relative to NEBD. White arrowheads highlight the Rpb3 foci after NEBD.
