## Supplementary material for "Distinct regulation across the transcription cycle drives mitotic transcriptional silencing": Figure S8

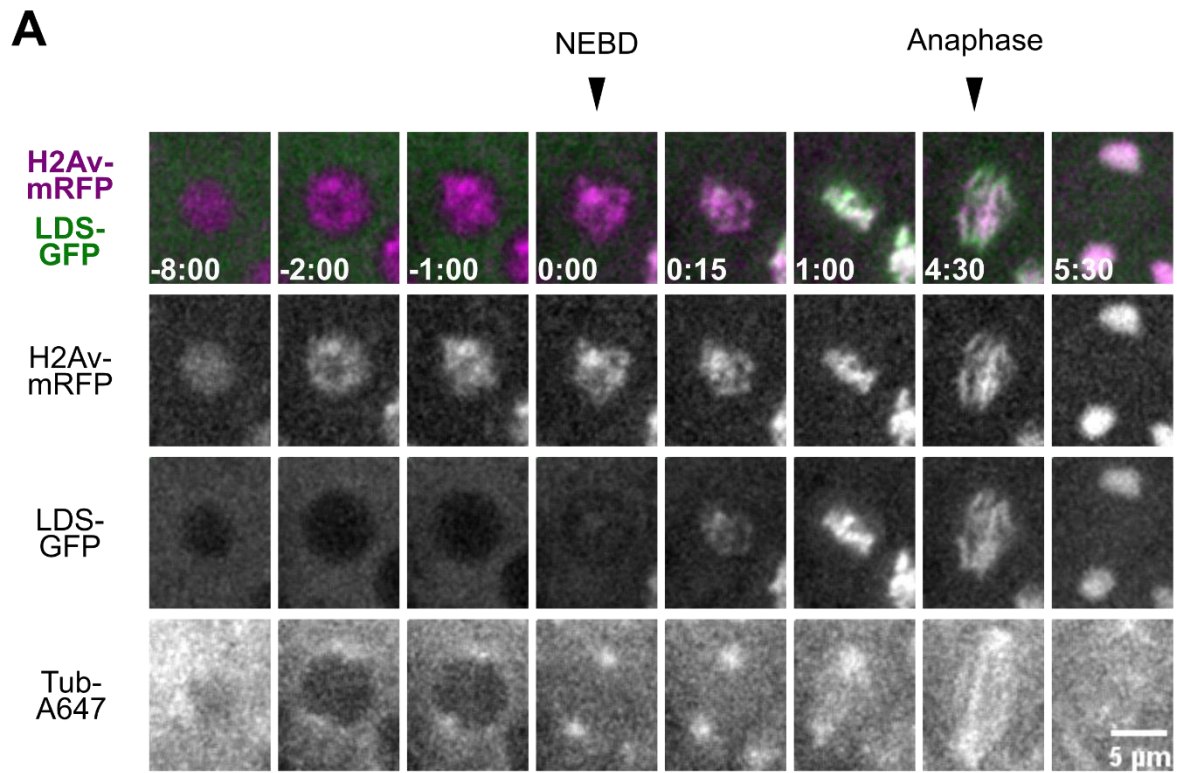

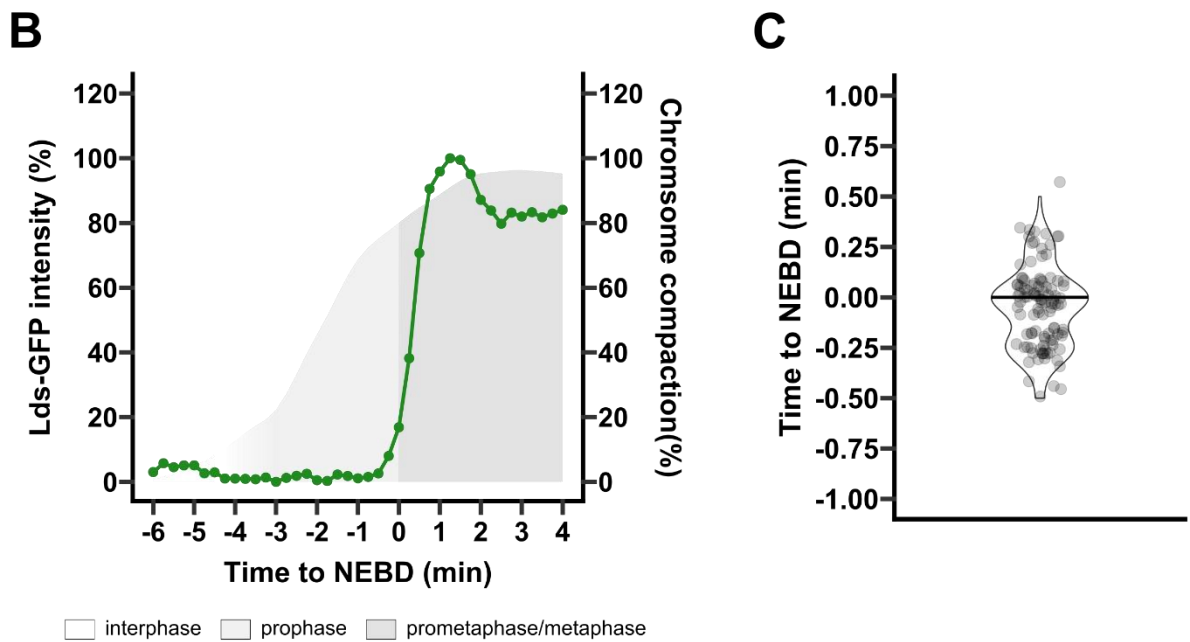

**Figure S8. Lds access chromatin at the time of NEBS.** A) Nucleus of a *Drosophila* syncytial embryo at nuclear cycle 12 expressing His2Av-mRFP (magenta), LDS-GFP (green), and injected with tubulin-Alexa647. Scale bar applies to all images. B) Quantification of LDS-GFP intensity in the chromatin area during cell cycle progression (green line). Grey area represents the progression of chromosome compaction colored according to the mitosis stages as per legend. C) Quantification of the first time frame in which Lds accesses the nuclei. N = 105 nuclei from 6 different embryos.
