## Supplementary material for "Distinct regulation across the transcription cycle drives mitotic transcriptional silencing": Figure S9

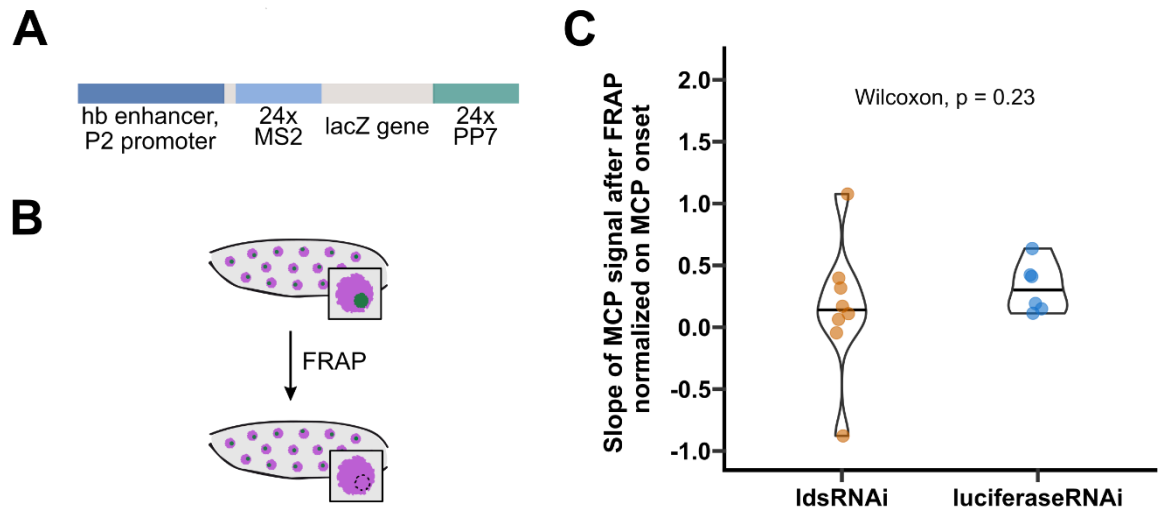

**Figure S9. The initiation rate is not affected in Lds-depleted embryos.** **A)** Scheme of the reporter used for this experiment. **B)** Experimental layout. Embryos in nuclear cycle 12, the MCP-GFP focus of one nucleus was photobleached. **C)** Slopes of the 5' transcription signal after FRAP in mitosis were normalized over the slope at the onset of transcription for each nucleus. No significant changes could be detected between the slopes in luciferase and lds RNAi embryos, as determined by the nonparametric Wilcoxon test (p-value in the panel).
