## Supplementary material for "Distinct regulation across the transcription cycle drives mitotic transcriptional silencing": Table S1

**Table S1. Theoretical intervals used to constrain parameter estimation in Figure 1.**

| **Table S1** | **Lower limit** | **Upper limit** |
| --- | --- | --- |
| $k_f$ (min<sup>-1</sup>) | 0.3<br>Calculated as the steepness necessary for the signal to decay from 0.9 to 0.1 in an entire cell cycle (15 min), according to the formula $k_f = 2\log(9)/\Delta t$, derived from the equation of initiation decay | 9<br>Calculated as the steepness necessary for the signal to decay from 0.9 to 0.1 in 2 time points (30 sec), according to the formula $k_f = 2\log(9)/\Delta t$, derived from the equation of initiation decay |
| $t_m$ (min) | -4<br>Average time of mitotic entry | 2<br>Average time of disappearance of the MCP signal |
| $\tau$ (min) | 0.5<br>Calculated as the time necessary to transcribe a reporter of 5.8 kb at a rate of 10 kb/min (fastest elongation rate reported in (Keller et al., 2023)) | 15<br>Length of a syncytial division, assumed as the longest time available to transcribe the reporter |
| $t_s$ (min) | -4<br>Average time of mitotic entry | 2<br>Average time of disappearance of the MCP signal |
| $\kappa$ (min<sup>-1</sup>) | 0.01<br>Small value close to zero | 9.2<br>Calculated as the coefficient necessary for a decay from 0.9 to 0.1 in two time points (15 sec), according to the relation $\kappa = 2.3/\Delta t$ derived from the premature termination term |

For each of the 93 nuclei fitted, at least 45 of the 50 restarts converged to the same final parameters (rounds that converged for each nucleus: $47.8 \pm 2.3$).
