## Supplementary material for "Distinct regulation across the transcription cycle drives mitotic transcriptional silencing": Table S2

**Table S2. Experimentally derived and theoretical intervals used to constrain parameter estimation.**

| **Table S2** | **Lower limit** | **Upper limit** |
| --- | --- | --- |
| $k_f$ (min<sup>-1</sup>) | 0.81<br>Calculated from the mean-SEM derived from the fitting of the 5' FRAP experiment in Figure 2G | 2.61<br>Calculated from the mean+SEM derived from the fitting of the 5' FRAP experiment in Figure 2G |
| $t_m$ (min) | -2.92<br>Calculated from the mean-SEM derived from the fitting of the 5' FRAP experiment in Figure 2G | -1.96<br>Calculated from the mean+SEM derived from the fitting of the 5' FRAP experiment in Figure 2G |
| $\tau$ (min) | 2.43<br>Calculated by dividing the reporter visible length (5.15 kb) by the elongation rate (first quartile of the distribution in Figure S6D) | 3.53<br>Calculated by dividing the reporter visible length (5.15 kb) by the elongation rate (third quartile of the distribution in Figure S6D) |
| $t_s$ (min) | -1<br>Earliest time at which the premature termination mechanism can act, according to the earliest time of no recovery in Figure S7A | variable<br>Last time point with detectable MCP-GFP signal on chromatin, different for each nucleus |
| $\kappa$ (min<sup>-1</sup>) | 0.01<br>Small value close to zero | 9.2<br>Calculated as the coefficient necessary for a decay from 0.9 to 0.1 in two time points (15 sec), according to the relation $\kappa = 2.3/\Delta t$ derived from the premature termination term |

For each of the 93 nuclei fitted, a mean of 39.2±13.8 restarts converged to the same estimated parameters.
