## Supplementary material for "Distinct regulation across the transcription cycle drives mitotic transcriptional silencing": Table S3

**Table S3. Theoretical intervals used to constrain parameter estimation for Lds- and Luciferase-depleted embryos.** For the parameters not reported, the intervals in Table S2 were used.

| Table S3 | Lower limit | Upper limit |
| --- | --- | --- |
| $t_s$ (min) | -0.07<br>First quartile of the $t_s$ distribution, as determined in Figure 4B | 2<br>Third quartile of the $t_s$ distribution, as determined in Figure 4B<br>To ensure consistency |
| $\kappa$ (min<sup>-1</sup>) | 0.01<br>Small value close to zero | 9.2<br>Calculated as the coefficient necessary for a decay from 0.9 to 0.1 in two time points (15 sec), according to the relation $\kappa = 2.3/\Delta t$ derived from the premature termination term |

For each of the 128 Luciferase RNAi nuclei fitted, a mean of $43.8 \pm 8.2$ restarts converged to the same parameters. For each of the 88 Lds nuclei fitted, $47.8 \pm 5.5$ restarts converged to the same estimated parameters.
