## Supplementary material for "Distinct regulation across the transcription cycle drives mitotic transcriptional silencing": Table S4

**Table S4. Fly strains used.**

| Genotype | Reference | Lab ID |
| --- | --- | --- |
| *w\**; *Rec* (*nos-MCP.EGFP*, *HistH2AvD-mRFP1*); *MKRS/TM6b* | This study, derived from (Garcia et al., 2013) (*nos-MCP.GFP*) and (Schuh et al., 2007) (*HistH2AvD-mRFP1*) | 1934 |
| *w[\*];[pbphi-hb proximal enhancer-P2 promoter-24x MS2-lacZ-24x PP7-αTubulin 3 UTR]* | (Fukaya et al., 2017) | 2000 |
| *y[1] w[\*]; P{hbP2-lacZ-MS2}JB38F;* | Bloomington BDSC # 60339 and (Garcia et al., 2013) | 1997 |
| *w[\*];[pbphi-hb proximal enhancer-P2 promoter-12x PP7-MS2-lacZ-αTubulin 3 UTR]* | (Fukaya et al., 2017) | 2003 |
| *w;* *Rec* (*nos-MCP.EGFP*, *HistH2AvD-mRFP1*); *G302-Gal4/MKRS* | This study, derived from (Araújo et al., 2022) (*G302-Gal4*) | 2005 |
| *His2Av-eBFP*, *PCP-mCherry*; *MCP-GFP* | (Fukaya et al., 2017) | 1923 |
| *y1 sc\* v1 sev21;P[y[+t7.7]v[+t1.8]=TRiP.HMS01389 \|attP2 (lds RNAi)* | Bloomington BDSC # 34980 | 1891 |
| *y[1] v[1]; P{y[+t7.7] v[+t1.8]=UAS-LUC.VALIUM10}attP2 (luciferase RNAi)* | Bloomington BDSC # 35788 | 2008 |
| *w; EGFP-Rpb3;* | (Cho et al., 2022) | 2006 |
| *w; PCNA-GFP;* | Gift from J. Raimundo; originally from (Blythe & Wieschaus, 2016) | 2009 |
| *w\**; *If/CyO*; *Lds-EGFP CRISPR [EGFP]*, *G302-Gal4/TM6b* | This study, derived from (Carmo et al., 2023) (*Lds-EGFP*) and (Pimenta-Marques et al., 2016) (*G302-Gal4*) | 1896 |
